# FUNGAL DYSBIOSIS CORRELATES WITH THE DEVELOPMENT OF TUMOUR-INDUCED CACHEXIA IN MICE

**DOI:** 10.1101/2020.06.29.171397

**Authors:** D.L. Jabes, Y.N.L.F. de Maria, D. Aciole Barbosa, K.B.N.H. Santos, L.M. Carvalho, A.C. Humberto, V.C. Alencar, R. Costa de Oliveira, M.L. Batista, F.B. Menegidio, L.R. Nunes

## Abstract

Cachexia (CC) is a devastating metabolic syndrome associated with a series of underlying diseases that greatly affects life quality and expectancy among cancer patients. Studies involving mouse models, in which CC was induced through inoculation with tumor cells, originally suggested the existence of a direct correlation between the development of this syndrome and changes in the relative proportions of several bacterial groups present in the digestive tract. However, these analyses have focus solely on the characterization of bacterial dysbiosis, ignoring the possible existence of changes in the relative populations of fungi, during the development of CC. Thus, the present study sought to expand such analyses, by characterizing changes that occur in the gut fungal population (*mycobiota*) of mice, during the development of cancer-induced cachexia. Our results confirm that cachectic animals display significant differences in their gut *mycobiota*, when compared to healthy controls. Moreover, identification of dysbiotic fungi showed remarkable consistency across successive levels of taxonomic hierarchy. Many of these fungi have also been associated with dysbioses observed in a series of gut inflammatory diseases, such as obesity, Colorectal Cancer (CRC), Myalgic Encephalomyelitis (ME) and Inflammatory Bowel Disease (IBD). Nonetheless, the CC-associated dysbiosis seems to be unique, presenting features observed in both obesity (reduced proportion of *Mucoromycota*) and CRC/ME/IBD (increased proportions of *Sordariomycetes, Saccharomycetaceae* and *Malassezia*). One species of *Mucoromycota* (*Rhyzopus oryzae*) stands out as a promising probiotic candidate in adjuvant therapies, aimed at treating and/or preventing the development of CC.

## 1.0 INTRODUCTION

Cachexia (CC) is recognized as a metabolic syndrome associated with several underlying diseases, such as cancer, chronic kidney disease and chronic heart disease, among others (1, 2). It is characterized by the reduction of muscle mass, depletion of body fat and generalized chronic inflammation (3). Its prevalence among patients with different types of cancer contributes to significantly decrease their life quality and expectancy, being an important cause of morbidity/mortality in more than 80% of advanced cancer cases and accounting for more than 20% of deaths (3,4,5). Until the 1980s, cachexia was attributed to anorexia and increased energy expenditure. However, enteral or para-enteral administration of nutritional supplements is not sufficient to reverse cachexia symptoms, refuting the hypothesis that nutrient deficiency is the main causative agent of this syndrome. During the late 1980s and early 1990s, cachexia came to be defined through a new concept, as a chronic inflammatory syndrome, and the metabolic changes associated with the cachectic condition derive from factors produced by both external agents (such as tumors), as well as by the affected organism (2).

Studies aimed at characterizing the microbiota of mice, during the development of cancer-induced cachexia, revealed significant alterations in the composition of their gut bacterial population (dysbiosis), when compared to healthy control animals (6,7,8,9,10,11). The main characteristic of such dysbiosis involves an increased proportion of *Enterobacteriaceae* in CC-affected animals – a phenomenon also observed during the development of several diseases and syndromes associated with gut inflammation, such as celiac disease (12), food allergy (13), gastric cancer (14), diabetes (15,16), obesity (17,18,19) and inflammatory bowel diseases (IBD), such as ulcerative colitis (UC) and Crohn’s disease (CD) (20), among others. More recently, cultivation of fecal material from cachectic mice in selective coliform media, identified *Klebsiella oxytoca* as the main representative of *Enterobacteriaceae* present in stool samples obtained from these animals (21). Based on this observation, it has been suggested that *K. oxytoca* could represent an intestinal pathobiont, capable of affecting both thickness and permeability of the mucus barrier that protects the digestive epithelium of mice, when over-represented in their gastrointestinal track (11,21,22).

These findings demonstrated that gut dysbiosis could be an important factor for the development of cancer-induced cachexia, raising the interesting possibility of developing adjuvant approaches to prevent/treat cachexia, by controlling the gut microbiota composition with the aid of specific antibiotics, prebiotics and probiotics. However, all CC studies carried out so far have focused solely on analyzing the bacterial microbiota, ignoring the possible existence of changes in the relative populations of fungi during the development of this syndrome. Thus, the present study sought to expand the scope of such analyses by characterizing the main alterations verified in the fungal population (*mycobiota*) present in the guts of mice, during the development of cancer-induced cachexia. In general, it was possible to verify the existence of significant differences involving the gut *mycobiota* of CC animals, in comparison to healthy controls. The fungal dysbiosis of the cachectic animals revealed alterations in the relative proportions of several fungal *taxa*, consistently distributed in successive levels of taxonomic hierarchy. In many cases, relative changes in these populations have also been described in dysbioses associated with other gut inflammatory diseases, although the CC-associated dysbiosis seems to display a unique signature. Moreover, some of the microorganisms found to be expanded among CC-affected animals display metabolic characteristics that may directly contribute to the development of cachexia. Finally, we identified fungal species preferably expanded in the gut of healthy animals, which can be grown axenically, under laboratory conditions. These latter microorganisms represent natural candidates to be tested as probiotics in adjuvant therapies, aiming at the treatment and/or prevention of cachexia.

## 2.0 MATERIALS AND METHODS

### 2.1. CACHEXIA INDUCTION IN C57BL/6 MICE AND COLLECTION OF STOOL SAMPLES FOR GUT *MYCOBIOTA* CHARACTERIZATION

The experiments described herein were conducted with 6-8-week old male mice, weighting 25-38 g. Animals were selected from colonies of C57BL/6 mice (Jackson Laboratory), which were bred and kept in plastic cages, in a rodent-only controlled environment, with low levels of external noise, 24 ± 1 °C average temperature and light/dark cycles every 12 hours. Food (Nuvilab CR1^®^ - Nuvital) and water were provided *ad libitum*. All animals were individually marked with identifying rings at the beginning of the experiment. A group of 10 animals was randomly chosen as a control group (CT), having received only subcutaneous injection with 200 μl saline solution in their right flank. The others (20) were similarly inoculated with 3.5×10^5^ LLC tumor cells (suspended in 200 μl of *Dulbecco’s modified Eagle’s medium*). Next, the animals were transferred to sterile cages and kept, under the conditions described above, for a period of 28 days, when cachexia symptoms could be verified (23,24). Stool samples were collected daily, from each mouse, until day 28. All samples were properly identified, frozen in liquid N_2_ and stored at −70°C. The genetic material obtained from the samples collected at day 28 was used to characterize the fecal microbiota of both CT and LLC-inoculated animals, as described below. All animals were sacrificed by decapitation, without anesthesia, immediately after collecting the last stool samples. Blood, organ and tissue samples from these mice were harvested and frozen at −70°C. This experimental design was evaluated and approved by the Ethics Committee for the Use of Animals in Research, at the University of Mogi das Cruzes (CEUA-UMC), under records n: 006/2013 and n: 009/2013.

### 2.2 ASSESSMENT OF CACHEXIA PROGRESSION

After inoculum, either with saline solution or LLC cells, all animals were weighed daily, on a precision scale (always at the same time), throughout the entire course of the experiment. This information was used to calculate the Cachexia Index (CI) in the LLC-inoculated animals, following the methodology described in 24. According to these calculations, 8 animals showed CI values ≥ 5, indicating full development of cachexia (24). These animals were further evaluated for their cachectic condition by histometric characterization of muscle (gastrocnemius) and adipose (epididymal) tissues (25), as well as by measuring the expression of molecular markers for muscular atrophy (*Atrogin*) and inflammation (*IL6 receptor*), by real-time quantitative PCR (qPCR) (see 26,27,28), for details). These 8 animals were designated Cachectic Group, or CC, which was further used for subsequent microbiome analyses. A similar group of 8 animals (the Saline Control Group, or SC) was selected among the CT animals, to establish a control group of similar size for future comparisons with the CC group.

### 2.3 SAMPLE PROCESSING AND DNA EXTRACTION

The extraction of fungal DNA from stool samples was performed with the aid of the *DNeasy Power Soil* kit, according to the manufacturer’s instructions (Qiagen). Integrity of the extracted material was evaluated through electrophoresis in a 1.5% agarose gel and purity was estimated through their A_260_//A_280_ ratios, using a NanoDrop 2000 spectrophotometer (Thermo Scientific). The final quantification of each DNA sample was determined with the aid of a Quantus fluorometer, according to the manufacturer’s specifications (Promega). The material from these extractions was used to characterize the fungal *mycobiota* by NGS sequencing, as described below.

### 2.4 PREPARATION AND SEQUENCING OF ITS1 *AMPLICON* LIBRARIES

Fungal populations were evaluated by comparing sequences from ribosomal internal spacer region 1 (ITS1), as described by 29. All *primers* used for library construction had, at their 5 ’ends, complementary regions to Illumina’s *Nextera XT* adapters. Thus, the *amplicons* obtained from each sample were subjected to a second *round* of PCR with *Nextera* adapters, adding specific *barcode* sequences at their ends. The entire process for preparing and purifying the *amplicon* libraries was performed as recommended by Illumina (https://support.illumina.com/documents/documentation/chemistry_documentation/16s/16s-metagenomic-library-prep-guide-15044223-b.pdf). Average sizes of the generated fragments were evaluated after electrophoresis in a *Bioanalyzer 2100* (Agilent), using the *High Sensitivity DNA Chip* and the libraries were quantified by qPCR, in an *ABI Prism 7500 Fast Sequence Detection System* (Applied Biosystems), with the aid of the universal quantification kit *NEBNext* (New England Biolabs), according to the manufacturer’s instructions. The final products of amplification were then aliquoted in a mixed sample of equimolar concentration and subjected to paired-end sequencing (2 × 300) in an *Illumina MiSeq* DNA sequencer, according to the manufacturer’s specifications (Illumina, Inc). Raw sequences obtained during the development of this study have been deposited at the Open Science Framework (https://osf.io/5fxgn) and SRA repository (https://www.ncbi.nlm.nih.gov/sra), under Provisional Submission ID # SUB7683387.

### 2.5 AMPLICON SEQUENCE ANALYSIS AND *MYCOBIOTA* CHARACTERIZATION

Sequence quality of individual reads was initially evaluated with the aid of *FastQC,* v 0.11. 9 (https://www.bioinformatics.babraham.ac.uk/projects/fastqc/) and *MultiQC,* v 1.7 (30), and subsequently analyzed through the *QIIME toolkit,* v 1.9.1 (31). During the pre-processing stage, individual *reads* were *demultiplexed,* paired and integrated into full size *amplicons*. Next, a filtering step removed all poor-quality amplicons from further analysis (filtered elements included a*mplicons* that displayed overlapping *reads* with less than 30 bases, strings with more than 10% divergence in the overlapping region, or that had more than three bases with quality index below Q20). Next, FASTA files from the selected *amplicons* were submitted to *USEARCH* v. 8 (32) to identify and eliminate chimeric artifacts, using the *UNITE* database, v. 8.0, as a reference (33,34).

During the processing stage, *Operational Taxonomic Units* (OTUs) were created, by clustering the individual sequences with a 97% identity threshold, with the aid of *SortMeRNA* (35) and *SumaClust* (36) software, using the *Open Reference* method. Taxonomic annotation was performed using the *RDP* classifier algorithm (37), with default settings, and the *UNITE* database v. 8 as reference (34). The resulting *OTU Table* was then filtered to remove contaminants (non-fungal, bacterial, plastid and archaea data) and *singletons*. Finally, a minimal abundance filter (A < 0.00001) was applied to remove OTUs underrepresented in the microorganism population. All filtering steps were performed using *QIIME* commands, as described by 38.

All these analytical steps have been implemented in a workflow described in a *Jupyter Notebook* file (39), in a Docker container, based on the *Dugong* project (40) and made available through *BioPortainer Workbench* (41). Details regarding this workflow can be found at: https://osf.io/5fxgn (DOI 10.17605/OSF.IO/5FXGN).

### 2.6 FUNGAL POPULATION AND DIVERSITY ANALYSES OF THE GUT *MYCOBIOTA* FROM SC AND CC MICE

Measurement of population ratios and Alpha/Beta diversity analyses were conducted with the aid of *MicrobiomeAnalyst* (42,43), using the *OTU Table* described above. To load the data, the *Low Count Filter* was adjusted to minimum values (*Minimum Count* = 0 and *Prevalence in Sample* = 10) and the *OTU Table* was rarefied by the number of sequences in the smallest library (*Rarefy to the Minimum Library Size*). All other parameters remained unchanged. Statistical significance of differences involving alpha diversity measurements between SC and CC groups was evaluated by a T-test (using p ≤ 0.05 as threshold). Variations in beta diversity were conducted with the aid of Bray-Curtis metrics (44) and visualized by a *Non-linear Multidimensional Scaling* (NMDS) algorithm, followed by statistical validation with PERMANOVA (using p ≤ 0.05 as threshold) (45). All these analyses were conducted using tools directly provided by *MicrobiomeAnalyst*. Dysbiosis Ratios (DR) were calculated and statistically validated as described in 46, using population ratios obtained from *MycrobiomeAnalyst*.

### 2.7 IDENTIFICATION OF DIFFERENTIALLY REPRESENTED TAXA IN SC AND CC MICE, USING LINEAR DISCRIMINATIVE ANALYSYS EFFECT SIZE (LEfSe)

Initially, core microbial populations were determined for each group separately (SC and CC), using the *QIIME* command *compute_core_microbiome.py* to select only OTUs present in ≥ 70% of the animals in the group. The *OTU Tables* containing the SC and CC *Core 70* microbiotas were then integrated, using the *QIIME merge_otu_tables.py* command, resulting in a *Core 70 OTU Table* for the entire SC X CC dataset. The use of 70% as a threshold to define *Core* microbiotas followed other studies described in the literature, where such limits vary between 50% and 80% (47,48,49,50,51,52,53,54,55,56). Next, the *Core 70 OTU Table* was submitted to the *QIIME summarize_taxa.py* script, integrating the results obtained for each taxon and converting their absolute prevalences into relative ratios. This relative *OTU Table* was then submitted to a *Linear Discriminant Analysis Effect Size* (LEfSe) evaluation, using the default settings of the *Galaxy LEfSe* tool, available at https://huttenhower.sph.harvard.edu/galaxy/.

## 3.0 RESULTS

### 3.1 CACHEXIA INDUCTION AND CHARACTERIZATION IN MICE

A group of 20 mice was submitted to LLC tumor cell transplantation and monitored for a 28-day period, under controlled conditions, as described in Materials and Methods. At the end of this period, animals were sacrificed and their weight gain was compared to a group of 10 control animals (CT group), to determine their respective Cachexia Indexes (CI) (Figure 1). Animals displaying CI ≥ 5 were further evaluated for other cachexia criteria, which included the observation of morphometric changes in adipose and muscle tissues (average decreases in adipocyte size and in the area occupied by muscle fibers, with consequent increase of interstitial muscular tissue), as well as increased expression of marker genes for muscular atrophy and inflammation (*Atrogin* and IL6R, respectively) (25,26,27,28) (Figure 1). In total, 8 LLC-injected animals displayed all characteristics of cachexia described above and these mice were selected for microbiome analyses, constituting the CC group. Accordingly, we selected a group of equal size among the CT animals to constitute the Saline Control (SC) group, which were used for further comparisons with the CC animals. Thus, stool samples obtained from SC and CC mice, at day 28, were further processed to characterize their fungal gut populations (*mycobiota*).

**Figure 1.**
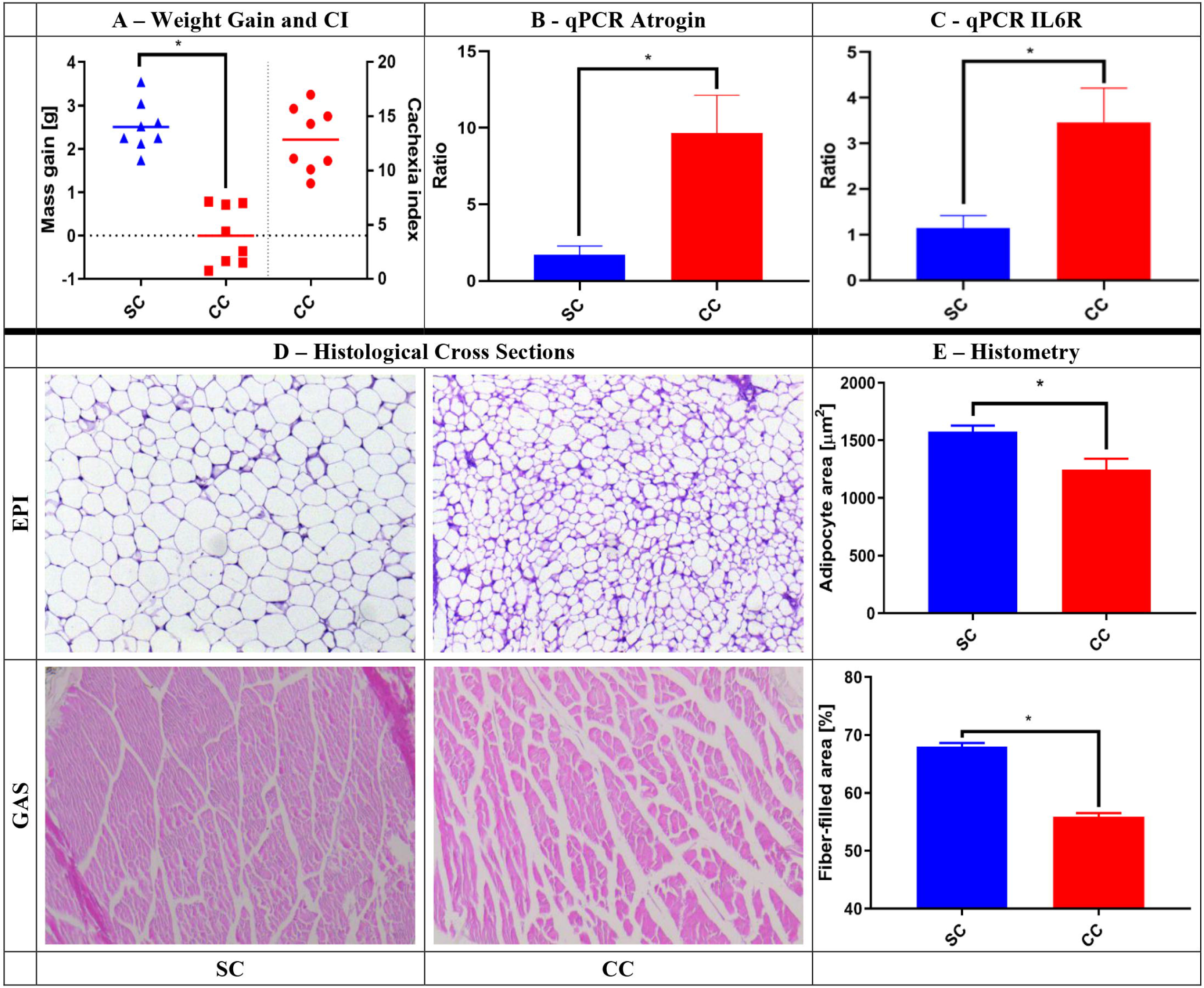
Characterization of cachexia in the animals selected for this study. A group of 16 animals was selected for the analyses described herein. Eight (8) of them received saline injection (SC group), while others were injected with LLC cells (CC group). After 28 days, the animals were evaluated for a series of characteristics typically associated with cachexia. Panel A shows the average weight gain of the SC and CC animals 28 days after injection (tumor weight subtracted), as well as the Cachexia Indexes (CI) among CC animals. Panels B and C show that the CC animals also showed increased relative expression of muscular atrophy and inflammation markers (Atrogin and IL6R, respectively), as verified by qPCR. Panel D shows examples of histological analyses made with epididymal adipose tissue (EPI) (top) and *gastrocnemius* muscle (GAS) (bottom) obtained from SC (left) and CC (right) animals, demonstrating significant reduction in the average adipocyte size and relative area of muscular fibers in CC animals. These reductions are confirmed by histometric evaluation of representative cross sections made with all animals from each group (Panel E). Magnification of adipose tissue: 20x; magnification of muscle tissue: 10x; * = p ≤ 0.05, after T test. Results for SC animals are shown in blue, while results for CC animals are shown in red.

### 3.2 CHARACTERIZATION AND COMPARISON OF FUNGAL GUT POPULATIONS (*MYCOBIOTA*) IN SC AND CC ANIMALS

*Amplicon* libraries referring to the ITS1 sequences were built for the animals of both SC and CC groups, as described in Materials and Methods and subjected to sequencing in an *Illumina MiSeq* DNA sequencer. The number of *amplicons* obtained for each animal, after all pre-processing steps, varied between 66,436 and 163,377, as shown in Supplementary Table 1. These sequences were submitted to an analytical pipeline, specially developed in our laboratory, generating a table of Operational Taxonomic Units (*OTU Table*) that was later used to analyze and compare the fungal gut populations (*mycobiota*) of SC and CC animals.

Surprisingly, initial analyses revealed that there was no alteration in fungal alpha-diversity when the fecal *mycobiota* of SC and CC animals were compared, since the absolute number of fungal species and OTUs did not show statistically significant differences (Figure 2A). Similar results were observed when using alternative alpha-diversity metrics (Shannon, Chao1, ACE, Simpson and Fisher) and/or after extending the analyses to other taxonomy levels (phylum to genus) (not shown). However, a beta diversity analysis revealed that the SC and CC animals display sufficient differences in the composition of their gut *mycobiota* to discriminate the two groups (Figure 2B). This analysis was performed with the aid of a *Non-linear Multidimensional Scaling* (NMDS) algorithm, using the Bray-Curtis index to compare the SC and CC *mycobiota* contents. The result from this analysis was subsequently evaluated by PERMANOVA (45), 2001), confirming that the SC and CC groups exhibit sufficient differences in the composition of their gut fungal populations to allow their discrimination at statistically significant levels (p = 0.045), although the two main components of the NMDS analysis do not provide visual separation between the two groups (Figure 2B).

**Figure 2.**
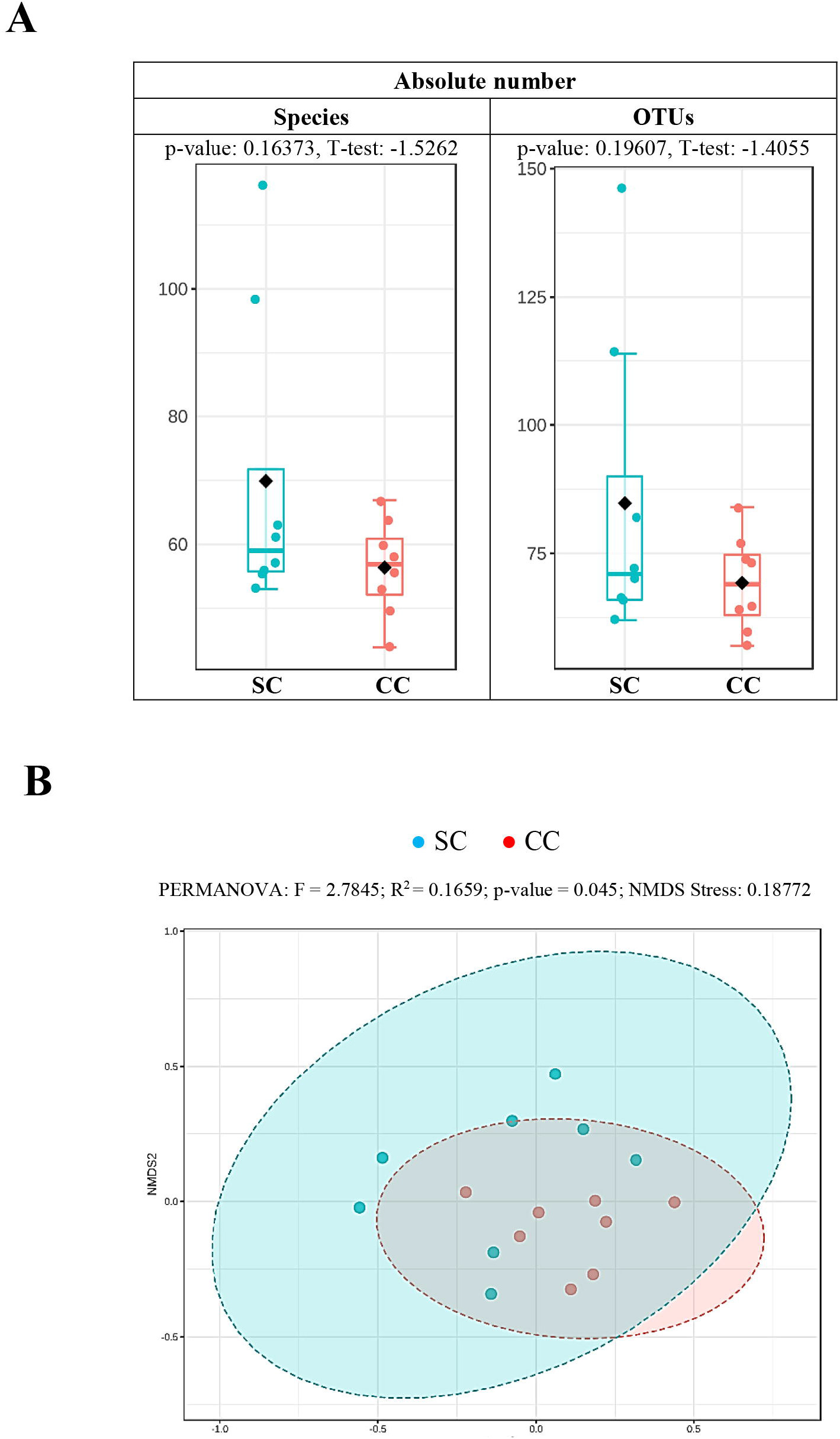
Alpha and Beta diversity analyses comparing the *mycobiota* of SC and CC animals. Panel A shows the results of alpha diversity analysis (absolute number of fungi). The *OTU Table* obtained from the sequences of ITS1 amplicons was filtered to keep only OTUs with abundance above 0.00001 and used to calculate the absolute number of fungi present in the stool samples from SC and CC animals, at different taxonomic levels. The figure shows the results obtained for the levels of species (left) and OTU (right). In both cases, it was not possible to verify significant changes in fungal alpha diversity between the two groups, since p-values obtained from such analyses, after T test, were always above 0.05. Similar results (not shown) were obtained with other alpha diversity metrics and/or after extending these analyses to additional taxonomy levels (see text for details). Panel B shows the result of a Beta-diversity analysis to evaluate differences in composition between the fungal populations present in the stool samples from SC and CC animals. This analysis was performed with the aid of a *Non-linear Multidimensional Scaling* (NMDS) algorithm (based on the Bray-Curtis index), using the same filtered *OTU Table* described above. The NMDS analysis shows that differences in *mycobiota* composition allow statistically significant discrimination between the SC and CC groups (at p = 0.045, as verified by PERMANOVA), although the two main components of the NMDS analysis do not allow us to visualize such distinction. Animals from the SC group are shown in blue, while animals from the CC group are shown in red.

As observed in Figure 3A, the vast majority of the fungal population in the gut of both SC and CC animals involves members from the phyla *Ascomycota* (~85-87%), *Basidiomycota* (~12-13%) and *Mucoromycota* (~0.4-1.5%). When the two groups are compared, however, it is possible to verify that CC animals display statistically significant reduction (~16X) in both *Mucoromycota*/*Ascomycota* and *Mucoromycota*/*Basidiomycota* ratios (Figure 3B-C). It is also possible to observe a slight increase (~1.7X) in the *Ascomycota/Basidiomycota* ratio of CC animals, although this difference did not display statistical significance under our experimental conditions (Figure 3D). Interestingly, similar population imbalances involving the main fungal phyla have also been observed in association with alternative gut inflammatory diseases and syndromes (see below). Unfortunately, it was not possible to detect statistically significant alterations involving other groups of fungi below the level of phylum, probably due to limitations involving the use of classical univariate statistical inference methods on microbiota compositional data (see 57).

**Figure 3.**
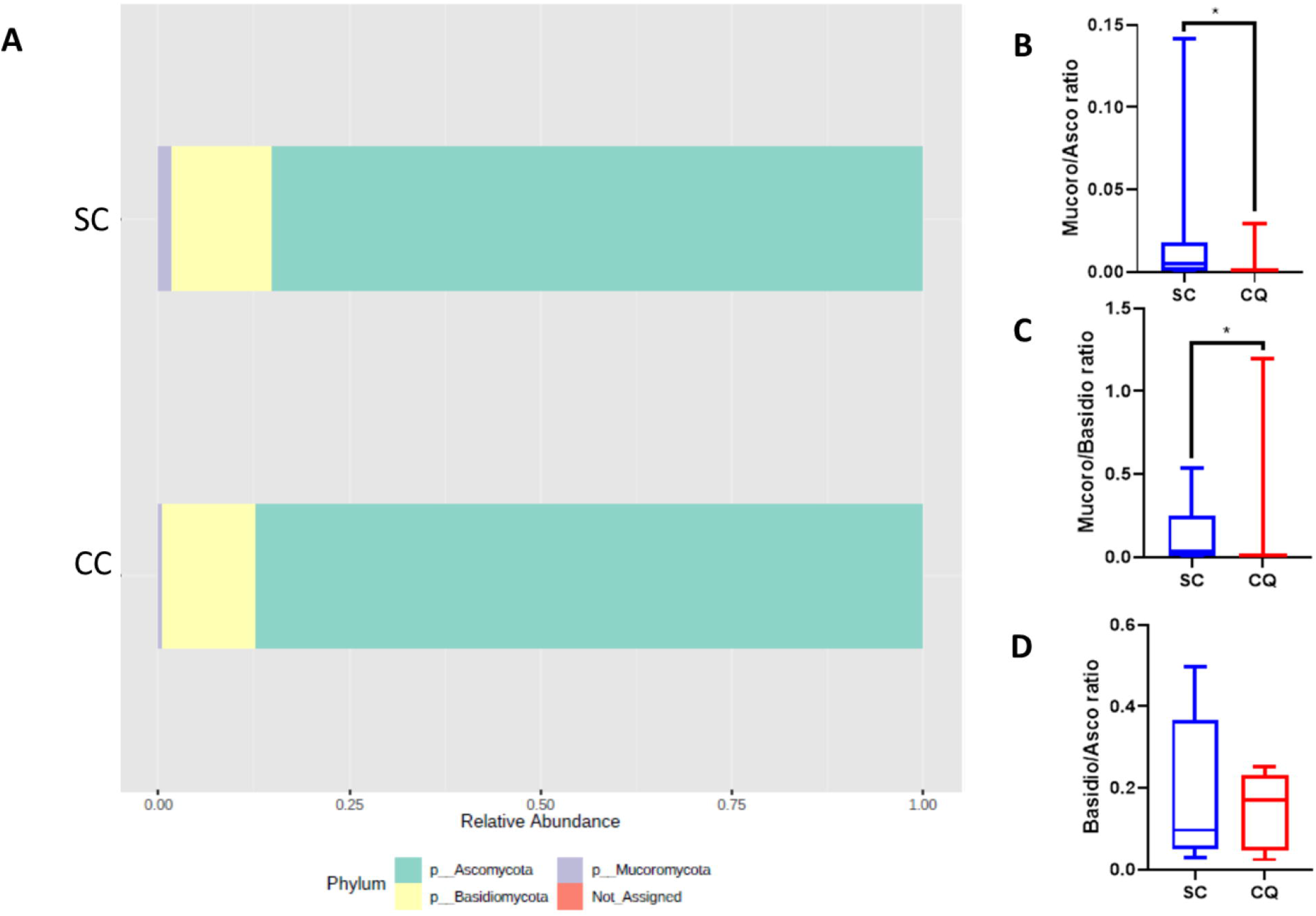
Population composition of the *mycobiota* from SC and CC animals and evaluation of fungal Dysbiosis Ratios (DR) associated with the development of cachexia. The *OTU Table* obtained from the sequences of ITS1 amplicons was filtered to keep only OTUs with abundance above 0.00001 and used to calculate the relative proportions of the main fungal phyla present in the gut of SC and CC animals. Panel A shows that the gut *mycobiota* of SC animals are mostly represented by *Ascomycota* (~85.1%), followed by *Basidiomycota* (~13.3%) and *Mucoromycota* (~1.5%). These proportions are shifted in the gut *mycobiota* of CC animals to ~87.2% (*Ascomycota*), ~12.5% (*Basidiomycota*) and ~0.4% (*Mucoromycota*). These values were used to calculate the Dysbiosis Ratios (DR) of SC and CC animals, involving the relative proportions of *Mucoromycota*/*Ascomycota* (Panel B), *Mucoromycota*/*Basidiomycota* (Panel C) and *Basidiomycota*/*Ascomycota* (Panel D). Results for SC animals are shown in blue, while results for CC animals are shown in red (* = p ≤ 0.05, after Mann-Whitney U test).

To overcome such limitations, the *mycobiota* of SC and CC animals were further compared through LEfSe (*Linear discriminant analysis Effect Size*), a widely used tool specifically developed to identify differentially represented elements from microbiome data (58) (Figure 4). This analysis was conducted with a *Core 70 OTU Table,* containing only OTUs present in, at least, 70% of the animals in each group. This strategy is often used to reduce effects related to individual variability among subjects, minimizing the effect of *taxa* that are over-represented in few individuals within a group, but not consistently present throughout the group (47,48,49,50,51,52,53,54,55,56). As expected, LEfSe analysis confirmed that members of *Mucoromycota* were present at higher proportion in the stool samples of SC animals, when compared to CC. Moreover, the SC-overrepresented *Mucoromycota* identified by LEfSe displayed remarkable consistency across all hierarchical levels of fungal taxonomy, involving representatives from the *Mucoromycetes* class*, Mucorales* order and *Rhizopodaceae* family, with emphasis on the species *Rhizopus arrhizus,* also classified as *R. oryzae* (59).

**Figure 4.**
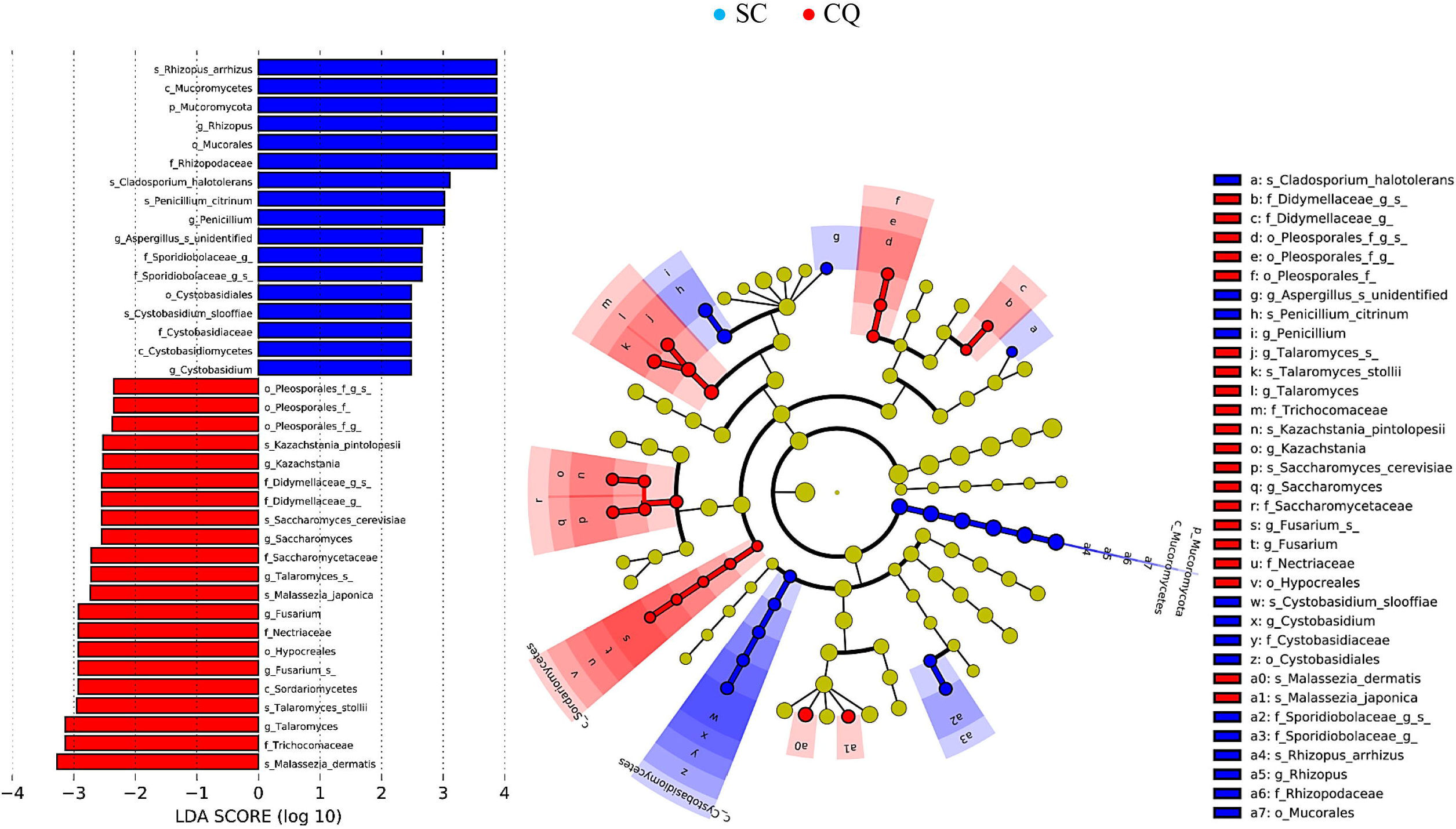
Identification of the main fungal taxa differentially represented in stool samples from SC and CC animals. The *OTU Table* obtained from the sequences of ITS1 amplicons was initially filtered to keep only OTUs with abundances above 0.00001 and present in at least 70% of the animals within each group, minimizing the effect of *taxa* over-represented in few individuals within a group, but not consistently present throughout the group. The *Core 70 OTU Table* derived from these filtration procedures was subjected to a LEfSe analysis, using p ≤ 0.05 and a modular limit for LDA Score ≥ 2 (−2 ≤ LDA Score ≥ 2) as stringency criteria to identify differentially represented taxa between CC and SC animals, respectively. All differentially represented taxa (at different taxonomic levels) are shown in the left panel of the figure and their distribution in a phylogenetic dendrogram is shown at the center panel. The concentric circles of the dendrogram show the taxonomic hierarchy, from phylum (innermost circle) to species (outermost circle) and the different microorganisms distributed in each node can be identified with the aid of the legend shown on the right panel. Microorganisms overrepresented in stool samples from SC animals are shown in blue, while microorganisms overrepresented in stool samples from CC animals are shown in red.

LEfSe also confirmed that there was no significant alteration in the overall distribution of *Ascomycota* and *Basidiomycota* phyla between SC and CC animals, but detected a series of fungal groups belonging to these two phyla that were differentially distributed in the stool samples from both SC and CC groups (Figure 4). Interestingly, the fungal groups that displayed the most consistent distribution across successive hierarchical levels of taxonomy highlight a series of taxa (mostly *Ascomycota*) that seem to be preferably expanded among CC animals (Figure 4). These include, for example, representatives from all taxonomic levels of the S*ordariomycetes* class, including members of the *Hypocreales* order and *Nectiriaceae* family, with emphasis on a species of *Fusarium, <u>w</u>*ith *<u>u</u>*nspecified *<u>c</u>*lassification [*wuc*]. Members of the *Saccharomycetaceae* family also showed population expansion in the stool samples of cachectic animals, including representatives of *Saccharomyces* and *Kazachstania* (with emphasis on *S. cerevisiae* and *K. pintolopesii*, respectively). Two representatives of *Tallaromyces* (*T. stollii* and another species [*wuc*]) were also identified as preferentially present in the feces of cachectic mice, when compared with SC animals. Additional *Ascomycota* displaying population expansion among CC animals involve representatives of the *Pleosporales* order, including members [*wuc*] of the *Didymellaceae* family. Interestingly, only two species of *Basidiomycota* were identified as over-represented in the feces of CC animals and both belong to the genus *Malassezia,* which congregates fungi traditionally involved with inflammatory processes, especially in the skin (*M. japonica* and *M. dermatis*). Animals from the SC group, on the other hand, presented expanded populations of *Basidiomycota* from the *Cystobasidiomycetes* class, including members of the *Cystobasidiales* order and *Cystobasidiomycetaceae* family, with emphasis on the species *Cystobasidium slooffiae*. Only three species of *Ascomycota* were identified as expanded among SC animals: *Cladosporium haloterians, Penicillium citrinum* and one species [*wuc*] of *Aspergillus*.

Finally, to test the reliability of these results, the relative proportions of some of the taxa described above were evaluated in SC and CC animals, using real-time quantitative PCR (qPCR). For this purpose, equimolar samples of DNA extracted from the stool samples from each mouse (the same DNAs used for the construction of *amplicon* libraries) were consolidated into two pools, representing the SC and CC groups. Samples of these pools were then used in qPCR experiments, with the aid of primers previously described in the literature as specific to the genera *Saccharomyces, Kazachstania* and *Malassezia,* as well as to the species *Saccharomyces cereviseae, Malassezia dermatis, Malassezia japonica, Rhizopus oryzeae* and *Penicillium citrinum* (60,61,62,63,64,65,66, 67). The *Ct* values obtained for each of these taxa were normalized by the *Ct* obtained with qPCR amplifications using the primer pairs used to build the fungal *amplicon* libraries and the relative quantification values, obtained for each taxon, are shown in Figure 5. These experiments confirmed the results obtained by LEfSe analysis, showing a significant expansion of representatives of *Saccharomyces* and *Kazachstania* (as well as the species *S. cereviseae*) within the gut of animals from the CC group, with more than one order of magnitude, in relation to SC. Representatives of the genus *Malassezia* (including *M. dermatis* and *M. japonica*) were also more represented in the pool of cachectic animals, although in smaller proportions (3 to 5X). Representatives of *Rhizopus oryzeae,* on the other hand, were detected exclusively in the pool of SC animals, which also corroborates the results obtained from LEfSe analysis. The only taxon that could not be amplified in these qPCR experiments was *Penicillium citrinum,* but we do not know if such negative results derive from under-representation of these fungi in the *mycobiota* from the two groups, or from problems regarding efficiency of the primers used in these experiments.

**Figure 5.**
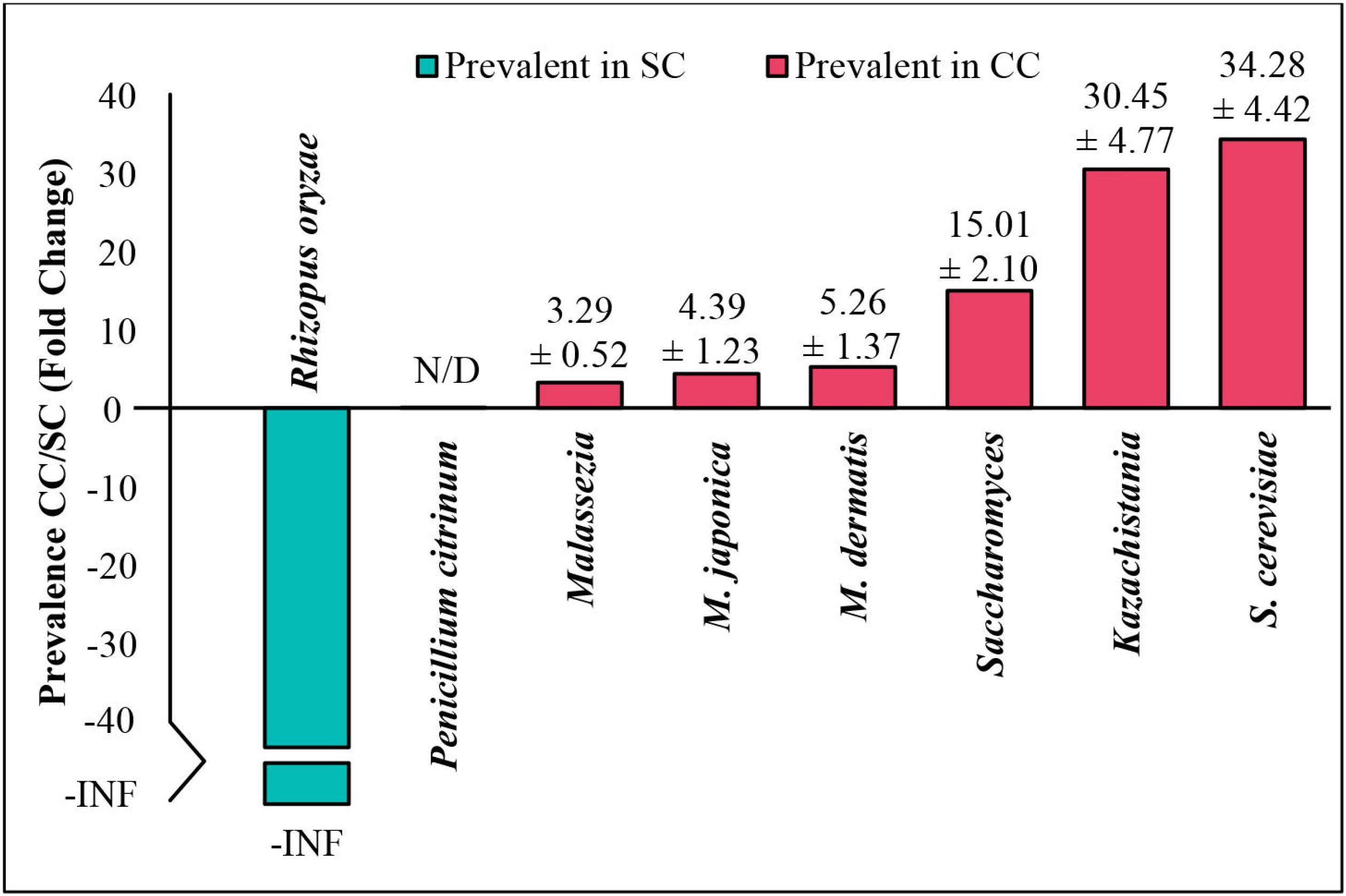
Relative quantification of different microorganisms in stool samples from SC and CC animals by qPCR. Equimolar samples of DNA extracted from stool samples obtained from each mouse were consolidated into two *pools,* representing SC and CC groups. Samples from these *pools* were then used in qPCR experiments, with primers previously described in the literature as specific to *Saccharomyceae, Kazachstania* and *Malassezia* genera, as well as to the species *Saccharomyces cereviseae, Malassezia dermatis, Malassezia japonica, Rhizopus oryzeae* and *Penicillium citrinum*. The *Ct* values obtained for each of these *taxa* were normalized by the *Ct* obtained after amplifying the pooled DNAs with the primer pair used to amplify the fungal ITS1 *amplicons*. Next, the normalized *Ct*s were used to calculate the relative prevalence of each genus/species in the CC *pool,* in relation to the SC *pool* (CC/CR). The relative quantification values for each taxon (*Fold Change*) are shown in the graph, as the mean ± standard deviation of three independent experiments (each performed in triplicate). The species *Rhizopus oryzae* was detected only in the DNA pool from SC animals, so its SC/CC ratio is represented in the graph as negative, to infinity (-INF). The species P*enicillium citrinum* was not detected (N/D) in any of the experiments, with either DNA *pool.*

## 4.0 DISCUSSION

Cachexia represents one of the most devastating clinical conditions that can occur with individuals affected by different types of cancer, since its high degree of morbidity results in significant reduction in the patients’ life quality. Moreover, treatments aimed at preventing or combating the development of CC often meet with limited success, since its onset and progression may be influenced by a variety of internal and external factors (2). In this sense, the existence of correlations between the development of cachexia and intestinal dysbiosis, as observed in mice, suggests that manipulation of gut microbial communities may assist in the development of novel adjuvant approaches to alleviate symptoms in cachectic patients, or to prevent its onset in individuals at risk of developing such syndrome. In this sense, the results presented herein demonstrate, for the first time, that intestinal dysbiosis associated with cachexia is not limited to changes in bacterial population, but also involves significant changes in the fungal population present in the gut of cachectic animals.

Analysis of relative ratios involving the main phyla of fungi are commonly performed to establish fungal *dysbiosis ratios* (DR), which have been proposed as quantitative indexes to describe alterations in *mycobiota* that accompany the progression of other dysbiosis-associated diseases (46,68). In this sense, the fungal DR observed in the gut of CC animals displays significant alterations in the relative ratios of both *Mucoromycota*/*Ascomycota* and *Mucoromycota*/*Basidiomycota* populations, confirming that CC animals display reduced proportions of *Mucoromycota* in their gut. Interestingly, a similar scenario has been reported by 69, who compared the gut *mycobiota* of lean and obese individuals. This similarity is noteworthy, since obesity and cachexia are both metabolic syndromes, directly associated with imbalances in energy metabolism, which affect both fat storages and muscular tissue. In the study conducted by 69, the relative reduction in the population of *Mucor* spp constituted the hallmark of the fungal dysbiosis verified in obese individuals. Thus, these authors suggested that the presence of chitosan in the cell wall of *Mucor* species could contribute to prevent obesity (a well-documented nutritional property of this polysaccharide) (70,71).

In this sense, *Rhyzopus oryzae*, the main *Mucoromycota* involved in the CC-associated dysbiosis described herein, may also act as a potential chitosan source in the gut of healthy animals, since this species has been employed in many industrial protocols aimed at chitosan production (72,73,74). This information raises the interesting possibility of testing chitosan and some of its derivatives, as food supplements, in adjuvant protocols aimed at treating or preventing cachexia, which may enhance the current pharmaceutical and biomedical applications of this polysaccharide (75). Additionally, *R. oryzae* has been reported to produce substantial amounts of various antioxidants and organic acids, including gallic acid (76). Interestingly, an extract of oil palm phenolics (OPP), containing 1,500 ppm gallic acid equivalent, has been shown to inhibit tumorigenesis, by mediating G1/S phase cell cycle arrest, delay inflammatory responses and attenuate cachexia symptoms in tumor-bearing mice (77,78). Gallic acid has also been shown to improve glucose tolerance and triglyceride concentration in obese mice (79) and suppress lipogenesis in humans, while concomitantly combating pro-inflammatory responses (80).

Alterations in relative ratios involving fungal phyla have also been detected and used to calculate DRs for a series of dysbiosis-associated diseases and syndromes, including Colorectal Cancer (CRC), Myalgic Encephalomyelitis (ME) (also known as Chronic Fatigue Syndrome, or CFS) and the two main forms of Inflammatory Bowel Disease (IBD): Crohn’s disease (CD) and Ulcerative Colitis (UC) (46,81,82,83,84). While the abovementioned diseases display alternative clinical manifestations, they all share common traits with obesity and cachexia, including increased permeability of the protective mucosal gut barrier and intestinal inflammation. Contrary to obesity and cachexia, however, CRC/ME/IDB/CD/UC do not affect adipose and muscular tissues and do not display gut dysbioses based on reduced ratios of *Mucoromycota*. Instead, these pathologies are all associated with fungal gut dysbioses characterized by altered *Ascomycota*/*Basidiomycota* ratios, when affected subjects are compared to their respective controls. As mentioned above, our analysis also indicated a small increase in the overall *Ascomycota*/*Basidiomycota* ratio, when CC animals are compared to SC, although this difference was not statistically significant, when evaluated by a classic Mann-Whitney U test. However, a more sophisticated comparison, performed with LEfSe, on the core *mycobiotas* of SC and CC animals, revealed that different groups of *Ascomycota* were preferentially expanded in the gut of cachectic mice – an information that was further confirmed by qPCR experiments, involving a series of representative *taxa*.

Interestingly, the main groups of *Ascomycota*, whose populations are expanded in CC animals, exhibit similar behavior in some of the diseases mentioned above. For example, a recent study aimed at evaluating the composition of the human intestinal *mycobiota*, demonstrated that members of the *Sordariomycetes* class (especially *Fusarium* and *Trichoderma*) are significantly enriched in rectal tissue samples from patients affected by CRC (85). Several species of *Fusarium* are recognized for causing infections involving both normal and immunosuppressed individuals and toxins produced by this group of fungi have already been shown to induce cell proliferation and affect different intestinal defense mechanisms (81). For example, *Fusarium* toxins can severely reduce thickness of the intestinal mucus layer, increasing its permeability and allowing microorganisms present in the gut lumen to reach the intestinal wall, triggering the production of immunoglobulins and pro-inflammatory cytokines (86). In fact, at least one *Fusarium* toxin, known as deoxynivalenol (DON) has already been correlated with the development of IBD (87), which is also in accord with the observations made by 82, who detected expansion of *Sordariomycetes* in the inflamed intestinal mucosa of CD patients.

Members of the *Saccharomycetaceae* family also showed increased population in the gut of patients affected by CD, with emphasis on *Candida glabrata,* a pathogen known to mediate inflammatory responses in the human intestine (82). In this sense, our studies point to a significant expansion involving the populations of two species of *Saccharomycetaceae,* among the CC animals: *Saccharomyces cereviseae* and *Kazachstania pintolopesii*. Moreover, the population expansion involving these two yeasts seems to occur at very high rates in the gut of CC animals (~30X), as verified by qPCR experiments. Interestingly, members of the *Kazachstania* genus show great phylogenetic proximity to *Candida* (88) and previous studies have even suggested that these two yeast genera could perform analogous functions in the intestinal micro-ecosystem of different animals (89). Paradoxically, studies by 82 identified an increased proportion of *Saccharomyces cerevisiae* in non-inflamed mucous membranes of healthy individuals, which led these authors to suggest that this yeast could play a beneficial role, preventing the development of CD. This proposal is in accord with results derived from other studies, involving both humans and mice, which suggested that *S. cerevisiae* could exert anti-inflammatory effects in host tissues, inducing the production of interleukin-10 (68,90).

However, 91 have argued otherwise, since the presence of *S. cerevisiae* exacerbated intestinal inflammation in a mice model of UC and increased the permeability of the protective intestinal mucosal barrier in these animals. In an independent study, 92 demonstrated that members of the *Saccharomycetaceae* family (particularly *S. cereviseae* and *Kazachstania unispora*) are able to trigger inflammatory responses through the interaction of α-mannans, present in their cell walls, with the lectin *dectin-2* of their host’s dendritic cells. The presence of *S. cerevisiae* also seems to potentiate the production of metabolites capable of exacerbating inflammatory processes in their hosts, including increased production and accumulation of uric acid (91). Finally, anti- *S. cerevisiae* antibodies (ASCAs) are often found in patients with CD and UC, being used as serological markers for differential diagnosis between these two forms of IBD (93,94). Taken together, these data reinforce the existence of a positive correlation between increased intestinal populations of *S. cereviseae* and *K. pintolopesii* with the development of gut inflammatory responses and increased permeability of the protective mucosal barrier, along the intestine (two characteristics also observed during the development of cachexia).

Other examples of *Ascomycota* displaying population expansion in the stool samples of CC animals have few references (or none) of association with the development of diseases. These fungi include members of the *Tricomaceae* family, with emphasis on the species *Talaromyces stollii,* as well as representatives [*wuc*] of the *Pleosporales* order and *Didymellaceae* family. Although there are no descriptions of dysbioses in the literature associating *T. stollii* with any type of disease, 95 detected a relative reduction in representatives of the genus *Talaromyces* in mice submitted to hypercaloric diets, while the species *Talaromyces islandicus* was positively correlated with the development of CRC (46). Members [*wuc*] of the *Pleosporales* order, however, have been associated with healthy mucosa in studies involving the occurrence of UV light-derived eye inflammation (96), as well as in the study on CD described by 82.

On the other hand, representatives of the genus *Malassezia,* the only *Basidiomycota* whose populations have been shown to expand in the gut of CC mice, have a long history of correlation with dysbioses associated with skin inflammation and dermatological diseases, since they are the main *mycobiota* components of the human and animal skin (97,98,99,100). Although the location of these fungi in the digestive tract has been previously described, the acceptance of these microorganisms as permanent inhabitants of the intestinal *mycobiota* is still a cause for debate (101). Nonetheless, 83 found a decrease in the relative population of *Malasseziales* in the intestine of patients affected by CD, while these same microorganisms were increased in the intestine of UC patients, suggesting that these fungi could be used as biomarkers to differentiate these two forms of IBD. More recently, 102 demonstrated that the species *M. rescricta* exacerbates symptoms of colitis in gnotobiotic mice and is present at a greater proportion in the colonic *mycobiota* of human subjects affected by CD, particularly in individuals with a specific polymorphism in the CARD9 gene. This gene encodes a protein responsible for recruiting immune cells to fight fungal infections and, curiously, patients affected by CD are known to develop anti-CARD9 antibodies (102). *Malassezia* representatives also compose a significant part of the microorganisms that invade the pancreatic duct, from the intestinal lumen, contributing to the development of Pancreatic Ductal Adenocarcinoma (PDA), through activation of the complement cascade, via interaction with mannan-binding lectins (MBLs) (103, 104). More recently, increases in the relative proportion of *Malassezia* have also been found in the gut of CRC patients and of children showing autoimmunity against pancreatic beta cells, which later developed type 1 diabetes (46,105).

The cachexia-associated fungal dysbiosis described herein is also characterized by additional fungal groups, whose relative proportions are decreased in CC animals (Figure 4). These include members of the *Cystobasidiomycetes* class (with emphasis on the species *Cystobasidium sloofiae*) and representatives [*wuc*] of the *Sporidiobolaceae* family. Although there are few reports regarding the involvement of these fungal groups with disease-associated dysbioses, increased populations of *Sporidiobolaceae* have been detected in cervical mucosa lesions from HPV-infected women at low risk of malignancy (106), as well as in the intestine of mice submitted to diets rich in animal proteins (107). Moreover, the relative abundance of *Cystobasidiomycetes* was positively correlated with adiposity, while negatively correlated with the serum concentration of LDL cholesterol in obese and lean subjects (69).

As mentioned above, obesity-associated fungal dysbiosis differs from the pattern observed in CRC/ME/IBD/UC/CD, as no significant changes have been detected in the relative ratio of *Ascomycota/Basidiomycota,* when obese individuals are compared to lean ones. In fact, the main characteristic found in obesity-associated *mycobiota* refers to a reduction in the proportion of *Mucoromycota,* a pattern also observed in our CC-associated dysbiosis (Figures 3 and 4). Thus, the CC-related dysbiosis seems to be unique, presenting features observed both in obesity (reduced proportion of *Mucoromycota*) and CRC/ME/IBD/UC/CD, with increased proportions of different groups of *Ascomycota* (such as *Sordariomycetes* and *Saccharomycetaceae*), as well as *Malassezia.*

Finally, it is worth mentioning that the population analysis shown in Figure 4 allowed identification of four fungal species preferably present in the gut of SC animals, which can be grown under axenic laboratory conditions (*Rhizopus oryzae, Cystobasidium sloofiae, Penicillium citrinum* and *Cladiosporium halotolerans*). These fungi are natural candidates to be tested (individually, or in combinations) for their eventual roles as probiotic agents, aiming at the prevention/treatment of cachexia (76,108). *Rhyzopus oryzae*, in particular, stands out as the most promising probiotic candidate, not only due to its capacity to produce bioactive substances, such as chitosan and gallic acid, but also to its historical use as a component of traditional oriental foods, which has granted it the status of a GRAS (Generally Recognized As Safe) filamentous fungus, according to the US Food and Drug Administration (FDA) (76).

## ACKNOWLEDGEMENTS

This study was financed in part by the São Paulo Research Foundation-FAPESP (http://www.fapesp.br) Grants #2017/13197-8 and 2017/08112-3, awarded to L.R.N and D.L.J. Y.N.L.F.M., F.B.M., D.A.B., K.B.N.H.S. and V.C.A. are recipients of scholarship grants from Coordenação de Aperfeiçoamento de Pessoal de Nível Superior – Brasil - CAPES (http://www.capes.gov.br/). L.M.C. and A.C.H are recipients of scholarship grants from Conselho Nacional de Desenvolvimento Científico e Tecnológico – Brasil – CNPq. Authors would like to thank the Core Facility for Scientific Research – University of São Paulo (CEFAP-USP/GENIAL [Genome Investigation and Analysis Laboratory]) for the Bioanalyzer 2100 services.

