## Supplementary Table 1 for "FUNGAL DYSBIOSIS CORRELATES WITH THE DEVELOPMENT OF TUMOUR-INDUCED CACHEXIA IN MICE"

### 8.0 SUPPLEMENTARY MATERIALS

**Supplementary Table 1 - Number of *amplicons* obtained after sequencing and pre-processing the ITS1 region of SC and CC animals**

| Animals | Total Number of Sequenced *Amplicons* |
|---|---|
| SC1 | 109,236 |
| SC2 | 66,436 |
| SC3 | 120,142 |
| SC4 | 135,404 |
| SC5 | 163,377 |
| SC6 | 144,145 |
| SC7 | 133,910 |
| SC8 | 153,817 |
| CC1 | 155,586 |
| CC2 | 141,331 |
| CC3 | 121,782 |
| CC4 | 124,370 |
| CC5 | 104,937 |
| CC6 | 137,693 |
| CC7 | 130,312 |
| CC8 | 120,530 |
| **Total** | **1,036,541** |
